# Spatiotemporal transformation of neural data reveals representations of erroneous behaviors

**DOI:** 10.64898/2026.07.04.736476

**Authors:** Duho Sihn, Sung-Phil Kim

## Abstract

Abnormal states such as erroneous behaviors are generally difficult to represent from neural data. However, such states are also known to have specific spatiotemporal features, indicating a feasibility of developing a method to focus on them. If a method can highlight these spatiotemporal features, it may effectively represent such abnormal states, helping evaluate abnormal brain functions. In the present study, we proposed the hierarchy of supported modules (HSM) to highlight spatiotemporal features that can represent abnormal states. HSM spatiotemporally transforms multidimensional neural time-series based on their spatiotemporal context. We evaluated HSM through decoding and similarity analyses using multiple publicly available datasets. In the HSM results, decoding accuracies were higher for erroneous behaviors than for normal behaviors, and similarities were lower between erroneous behaviors and normal behaviors than between normal behaviors, demonstrating the ability of HSM to capture the spatiotemporal features of erroneous behaviors. Surprisingly, many parts of these results were also present even before HSM learning, showing the virtue of HSM as a simple-to-use method. The proposed HSM method may help elucidate the mechanisms underlying erroneous behaviors.

## 1. Introduction

The human brain has evolved to optimize human performance. As such, the human brain is unlikely to represent suboptimal states such as erroneous behavior. This may lead to a lack of neural representations for erroneous behaviors in the brain. However, the spatiotemporal structures of neural activity during erroneous behaviors are not merely random, but known to contain representations of these states. For example, neural traveling waves in the brain, which consist of sequences of neural activities, propagate in opposite directions during erroneous behaviors (Alamia et al. 2023). When a human commits errors, there are characteristic event-related potentials in mid-frontal area (Holmes and Pizzagalli, 2008). Furthermore, prolonged blood-oxygen-level-dependent (BOLD) responses are observed during erroneous behaviors (Janik et al., 2022). To explore neural representations during abnormal states, it is therefore important to consider the spatiotemporal context of neural activity, beyond a single pattern of neural activity.

A single pattern of neural activity does not have one-to-one correspondence to a single piece of neural information. This is well demonstrated by the phenomenon of trial-to-trial neural variability in which neural activity patterns vary over trials in response to the same neural information (Nogueira et al., 2020; Zhang et al., 2022; Nakuci et al., 2023). This variability emerges as specific spatiotemporal structures that are distinct from noise (Ni et al., 2022; Ribeiro et al., 2024; Akella et al., 2025). Previous model studies have shown that recurrently-networked structures can yield structured variability (Kotekal and MacLean, 2020; Negrón et al., 2024), which has also been verified experimentally (Xiao et al., 2024). In networked neural systems, it is more important to focus on the spatiotemporal context of neural activity than inspecting instantaneous spatial neural activity patterns alone (for a review, see Ju and Bassett, 2020). If an identical pattern of neural activity in different spatiotemporal contexts can be disentangled, a more accurate assessment of neural information dynamics can be made. Therefore, it is important to disentangle, or more generally transform, neural data by taking spatiotemporal context into account. This may improve our understanding of representations of abnormal states.

In relation to neural variability, a method for disentangling neural data to reduce independent noise has been introduced (Lecoq et al., 2021). However, this method did not address the spatiotemporal context of neural data. Because the recurrently networked structures of the brain generate the spatiotemporal context of neural data, recurrent neural networks can be used to measure this spatiotemporal context (Durstewitz et al., 2023). One such example is the estimation of the dynamical system of neural data through recurrent neural networks (Koppe et al., 2019). However, recurrent neural networks in such studies are trained solely to describe dynamical systems accurately. In the case of disentangling task-related neural data, previous work has focused on stimulus-feature disentangling rather than neural-activity disentangling (Dado et al., 2024). In the case of more general neural data transformation, previous studies have mainly focused on transformations from spatiotemporal space to another space (Shaeri and Sodagar, 2023). Transformations from one spatiotemporal space to another spatiotemporal space conditioned on spatiotemporal context have been less explored.

One possible way of transforming neural data based on spatiotemporal context is to *support* the current pattern of neural data by spatiotemporal generation of future data from past patterns of neural data within a neural-network architecture. If the spatiotemporal contexts of past patterns of neural data differ, then the same current pattern of neural data will be supported differently, thereby disentangling neural data. Moreover, although current patterns of neural data may differ, appropriate support from past histories may transform them into similar representations.

In general, neural data are costly in the sense that recording them over long durations is difficult. This leads to limited sample sizes, which in turn limit the size of neural networks. The required sample size increases explosively as the size of neural networks increases (Ehrenfeucht et al., 1989; Maass, 1994). One way to overcome this problem is to use relatively small modules that engage only a subset of specific samples. A representative example of this type is a mixture of experts (Jacobs et al., 1991); moreover, the brain also has such modular structures (Meunier et al., 2010; O’Doherty et al., 2021). A mixture of experts is well suited to well-clustered samples (Chen et al., 2022); however, neural data are not generally well clustered. To resolve this problem, we devised a hierarchical structure of modules with a gradient of participation. At the lower hierarchy, many modules participate simultaneously in computation so as to address the problem of non-clustered neural data. At the higher hierarchy, a smaller number of modules participate in computation so as to overcome the problem of limited sample size.

Unlike in a mixture of experts, the modules participating in computation are determined by the modules themselves. In each module, the current pattern of activity is supported by spatiotemporal generations of future data from past patterns of activity; correctly supported modules participate in computation. Modules engaged solely in specific neural data should process that neural data while reflecting its entire spatiotemporal context. This leads us to make modules in higher hierarchies supported by longer past patterns of activity.

In this study, we referred to this neural-network scheme of modules as the hierarchy of supported modules (HSM). First, we examined whether HSM transforms (disentangles) synthetic data based on their spatiotemporal context for a ground-truth test. We then applied HSM to various publicly available neural datasets: human electroencephalogram (EEG) data and human functional magnetic resonance imaging (fMRI) data. Through analyses of these neural data, we investigated the effects of neural data transformation.

## 2. Materials and Methods

### 2.1. Hierarchy of supported modules (HSM)

The basic structure of HSM was as follows. HSM has a hierarchical structure and consists of hierarchies *h* = 1, 2, …. Each hierarchy *h* consists of several modules *m* = 1, 2, …. Each module *m* consists of several units *i* = 1, 2, …, which are artificial neurons in neural networks.

To define HSM, we first introduced a novel neural-network activation function, namely the variation-controlled activation function (VCAF). VCAF was devised to fix the level of variation in unit activity. Let *x*_*i*_ be the activity of unit *i* in module *m*, and let 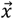 be the activity vector of the units. By applying VCAF to 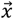, the activity of the units is transformed into:

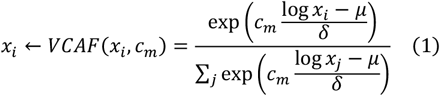

where *μ* is the average of log *x*_*i*_ over *i, δ* is the average of |log *x*_*i*_ − *μ*| over *i*, and *c*_*m*_ is the control parameter of module *m*. The vector 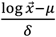 represents a standardized vector with zero mean and unit variation, so the variation of *VCAF*(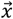, *c*_*m*_) is completely controlled by parameter *c*_*m*_. If *c*_*m*_ is fixed, the variation of *VCAF*(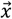, *c*_*m*_) is invariant to changes in 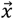, which is why the function is “variation-controlled.” A large *c*_*m*_ increases the variation of 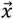. Note that a unit activity *x*_*i*_ is always non-negative and has an average value of 1, because it is transformed by VCAF.

A “supported” module means that the activity of its units is supported by activity history. Specifically, the activity of unit *i* of module *m* in hierarchy *h* at time *t, x*_*i,m,h*_(*t, c*_*m*_) under control parameter *c*_*m*_, is:

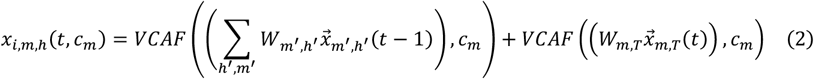

where *W*_*m*_′_,*h*_′ is the inter-hierarchy weight matrix from (*m*^′^, *h*^′^) to (*m, h*), 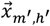 (*t* − 1) is the activity vector of the units in module *m*^′^ in hierarchy *h*^′^ at time *t* − 1, *W*_*m,T*_ is the weight matrix from the history 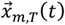 to the current activity 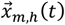, and 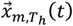 is the history of 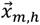 from time *t* − 1 to time *T*_*h*_ . Note that the weight matrices were non-negative and were magnitude-normalized by the sums of their column and row vectors. The duration *T*_*h*_ was longer in the upper hierarchy, representing the longer time scale. The first term represents external inputs, and the second term represents the “supported module,” which is supported by the internal history.

Note that a small *c*_*m*_ renders 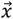 a nearly trivial vector with value 1, so the effects of inputs are negligible, whereas a large *c*_*m*_ highlights a few points in 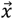 so that it becomes an informative transmission from the inputs. In other words, only modules with large *c*_*m*_ are effective.

Let *μ*_*m*_ be the average of log *x*_*i,m,h*_(*t, c*_*m*_) over *i*, and let *δ*_*m*_ be the average of |log *x*_*i,m,h*_(*t, c*_*m*_) − *μ*_*m*_| over *i*. A large *δ*_*m*_ indicates that the unit activities in module *m* are informative. The variation of *δ*_*m*_ in hierarchy *h*, namely 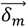, was controlled by the control parameter *C* of hierarchy *h*:

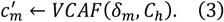

Here, 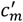 was the variable determined at each time based on *C*_*h*_, and *C*_*h*_ was the pre-fixed value, set to be larger in the upper hierarchy, thereby guaranteeing the large variation of 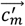. A large value of *C*_*h*_ emphasizes the retention of only a few highly informative modules, producing a strong “mixture of experts” effect. These transformed 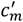 at time *t* were re-applied to the calculation of *x*_*i,m,h*_(*t, c*_*m*_), thereby re-calculating the final value of 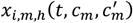 for each time *t*:

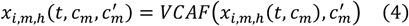

where only modules with large 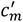 are effective in hierarchy *h*. This brings about the effect of “a mixture of experts.”

In this study, the hierarchy height was 3. The number of modules in each hierarchy was 16, and the number of units in each module was 16. The durations *T*_*h*_ of internal history at each hierarchy were 8, 16, and 24 for the 1st, 2nd, and 3rd hierarchies, respectively. All units in adjacent hierarchies were fully connected, and these connections were initialized with uniform random numbers. We also devised the virtual data hierarchy, namely *h* = 0, which is connected from *h* = 1, representing the neural time-series data from the unit activities in *h* = 1. HSM was learned by the Hebbian rule (Figure 1).

**Figure 1.**
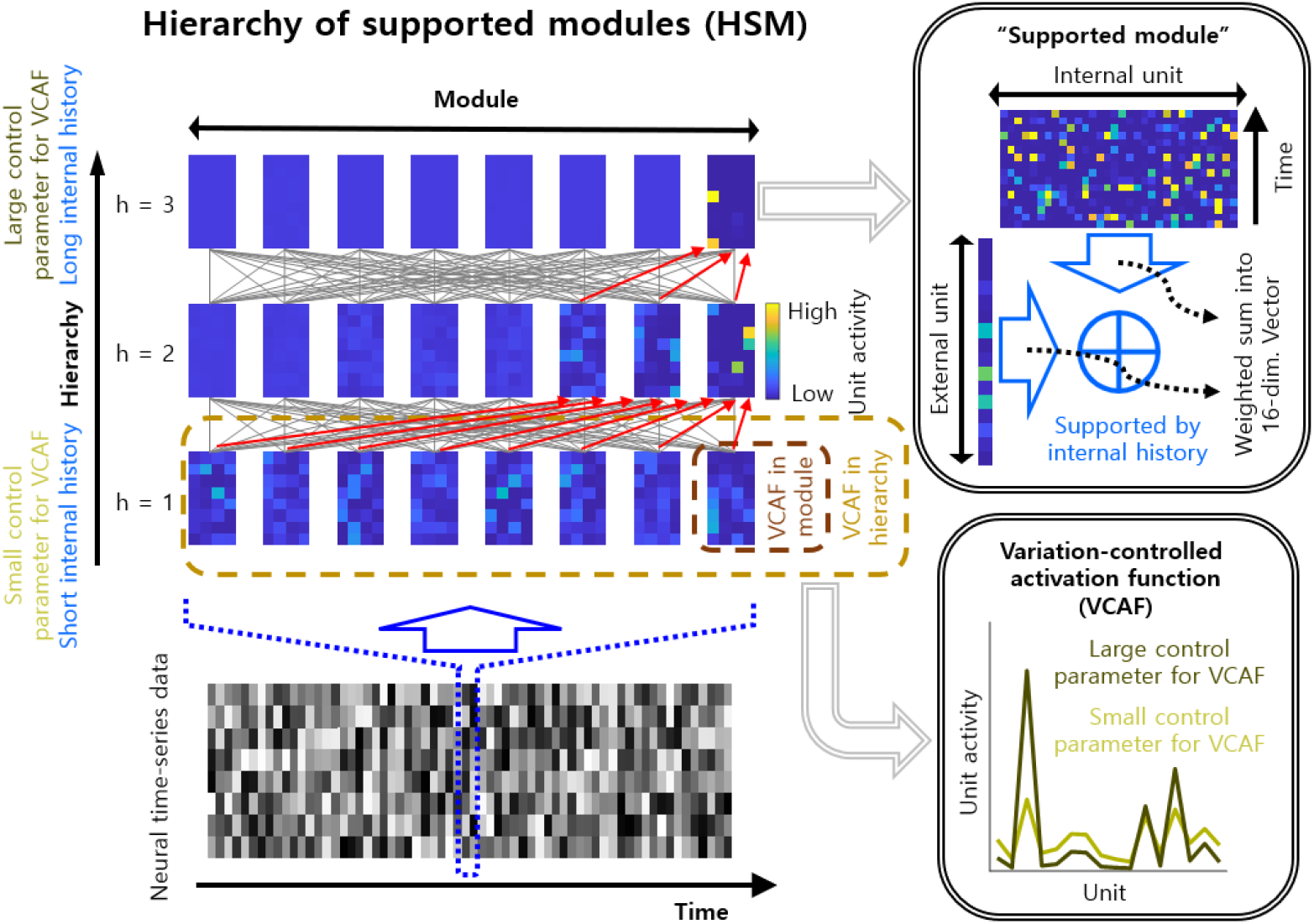
Hierarchy of supported modules (HSM). HSM has a hierarchical structure and consists of hierarchies *h* = 1, 2, 3. Each hierarchy *h* consists of several modules *m* = 1, 2, …. Each module *m* consists of several units *i* = 1, 2, …, which are artificial neurons in neural networks. Each module is supported by its internal history, namely the “supported module,” transforming (disentangling) unit activities based on different histories. The duration of internal history was longer in the upper hierarchy, representing the longer time scale. The variation-controlled activation function (VCAF) sets the variation of unit activities in the module to the intended level. The variation in the module was controlled by VCAF in the hierarchy. The control parameter for VCAF in the hierarchy was larger in the upper hierarchy, producing a stronger “mixture of experts” effect. We also devised the virtual data hierarchy, namely *h* = 0, which is connected from *h* = 1, representing the neural time-series data from the unit activities in *h* = 1.

#### 2.1.1. Details of HSM

Connections between units in adjacent hierarchies were bidirectional. However, connections in the bottom-up direction were eight times stronger than those in the top-down direction, making HSM primarily a tool for data representation.

The control parameter *c*_*m*_ of module *m* in hierarchy *h* was initially set to *h*/3. The control parameter *C*_*h*_ of hierarchy *h* was set to *h*/3. Then, the last control parameter 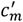 was calculated based on *C*_*h*_. If 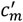 exceeds 2, 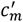 was reset to 2.

HSM was learned by the Hebbian rule. Let *w*_*ij*_ be the weight from unit *j* to unit *i*. If both activities 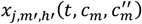 and 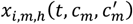 of units *j* and *i* exceed 3, the weight *W*_*ij*_ was updated to 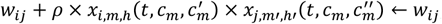, where the learning rate was *ρ* = 10^−4^ × 256^−1^. The weight matrices were magnitude-normalized by the sums of their column and row vectors once every 240 time steps. Unfortunately, this learning tended ultimately to lead to trivial solutions such as temporally unchanged unit activities. To suppress this, we set *w*_*ij*_ /240 ← *w*_*ij*_ when the averages of the activities of units *j* and *i* over the recent 240 time steps exceeded 1.5.

Rapidly changing unit activity and variation control could make HSM learning difficult. To smooth unit activity over time steps, we set 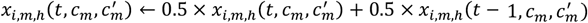. Let *μ*_*m*_(*t*) be the average of log *x*_*i,m,h*_(*t, c*_*m*_) at time step *t* over *i, δ*_*m*_(*t*) be the average of |log *x*_*i,m,h*_(*t, c*_*m*_) − *μ*_*m*_(*t*)| at time step *t* over *i*, and let *μ*_*h*_(*t*) be the average of log *δ*_*m*_(*t*) at time step *t* over *m* in hierarchy *h*. Let *δ*_*h*_(*t*) be the average of |log *δ*_*m*_(*t*) − *μ*_*h*_(*t*)| at time step *t* over *m* in hierarchy *h*. To smooth variation control over time steps, we set *δ*_*h*_(*t*) ← 0.5 × *δ*_*h*_(*t*) + 0.5 × *δ*_*h*_(*t* − 1). Smoothing unit activity and variation control in this way may facilitate HSM learning.

Conversely, unit activity and variation control that change too slowly will prevent HSM from properly reflecting changes in neural data. Let *I*_*τ*_(*t* − *s*) be a characteristic function such that *I*_*τ*_(*t* − *s*) = 1, if *s* ≤ *τ*, and *I*_*τ*_(*t* − *s*) = 0, if *s* > *τ*. Let *J*_*y*>*c*_(*t* − *s*) be a characteristic function such that *J*_*y*>*c*_(*t* − *s*) = 1, if *y*(*t* − *s*) > *c*, and *J*_*y*>*c*_(*t* − *s*) = 0, if *y*(*t* − *s*) ≤ *c*. To prevent persistent unit activity, 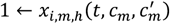, if 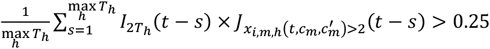. To prevent persistent variation control, 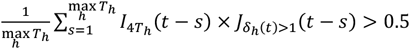. Breaking unit activity and variation control in this way may allow HSM to reflect changes in neural data properly.

### 2.2. The control model for HSM

To compare HSM with a control model in synthetic and real data analysis, we designed the model that has same hierarchy as HSM but is not supported history, as the control model. The control model had the same number of hierarchies and units as HSM: h = 1, 2, and 3, and 256 (16 × 16) units in each hierarchy. Moreover, the control model had same control parameter as HSM; The control parameter *C*_*h*_ of hierarchy *h* was set to *h*/3. However, the control model lacked the modular structure of HSM, as well as the supporting via spatiotemporal history.

### 2.3. Synthetic data

The synthetic data consisted of images of a unimodal hill. The peak of this hill moved vertically or horizontally and then returned, forming a movie that represents multidimensional time-series data. Each trajectory consisted of 16 time steps of peak movement, 8 time steps for going and 8 time steps for returning. There were 8 vertical trajectories and 8 horizontal trajectories. In the movie of peak movements, adjacent trajectories were displayed consecutively. All 16 trajectories of 16 × 16 = 256 time steps constituted one set of displays. The whole movie consisted of 40 repeated sets of displays with 256 × 40 = 10240 time steps. For learning, the whole movie was exposed to HSM 10 times with 10240 × 10 = 102400 time steps.

### 2.4. Publicly available datasets

#### 2.4.1. Human EEG dataset recorded during the gameplay task

To study representations of erroneous behaviors, we utilized a publicly available dataset (Cavanagh and Castellanos, 2021). This dataset was available at: https://openneuro.org/datasets/ds003517/versions/1.1.0. This EEG dataset was recorded using a 63-channel EEG system (actiCHamp, Brain Products GmbH, Germany) at a sampling rate of 500 Hz during a gameplay task. The EEG data were recorded from 17 healthy participants (6 females), and the mean age of the participants was 29.94 years (standard deviation = 5.02). Participants played the 8-bit video game for about 45 minutes. During gameplay, participants operated a spacecraft, performing evasive maneuvers to avoid obstacles and launching missiles at enemy spacecraft, yielding six behavioral events: launching missiles (Shooting 1), hitting an enemy with a missile (Shooting 2), collecting health (Collecting 1), collecting missiles (Collecting 2), crashing into an enemy (Crashing), and crashing into a wall (Error) (Cavanagh and Castellanos, 2016).

To fit the fixed duration *T*_*h*_ of internal history in HSM, the dataset was downsampled to 10 Hz just before exposure to HSM. The dataset was exposed to HSM 5 times.

#### 2.4.2. Human fMRI dataset recorded during the working memory task

To study representations of erroneous behaviors, we utilized a publicly available dataset (Braver et al., 2025). This dataset was available at: https://openneuro.org/datasets/ds003465/versions/1.0.7. This fMRI dataset was recorded using a 3 T magnetic resonance imaging (MRI) system (Prisma, Siemens, Germany) with a repetition time (TR) of 1.2 seconds during a working memory task. The fMRI data were recorded from 55 healthy participants (34 females), and the mean age of the participants was 31.7 years (standard deviation = 5.9). Participants performed the Sternberg task as a working memory task for about 24 minutes. During the Sternberg task, participants temporarily remembered a list of words and then responded with the remembered word after a few seconds, yielding four types of trials: correctly responded trials with low cognitive demand type 1 (Low demand 1), correctly responded trials with low cognitive demand type 2 (Low demand 2), correctly responded trials with high cognitive demand (High demand), and incorrectly responded trials (Error) (Braver et al., 2021; Etzel et al., 2022).

The dataset was exposed to HSM 120 times.

#### 2.4.3. Human EEG dataset recorded during the inhibition task from patients with Parkinson’s disease

To study representations of erroneous behaviors and brain diseases, we utilized a publicly available dataset (Singh et al., 2023a). This dataset was available at: https://openneuro.org/datasets/ds004580/versions/1.0.0. This EEG dataset was recorded using a 63-channel EEG system (BrainVision, Brain Products GmbH, Germany) at a sampling rate of 500 Hz during an inhibition task. The EEG data were recorded from 48 healthy controls (HC; 22 females) and 99 patients with Parkinson’s disease (PD; 32 females), and the mean ages of HC and PD were 71.13 and 68.55 years, respectively (standard deviation = 7.56 and 8.10). PD was determined by the United Kingdom PD Society Brain Bank criteria. Participants performed the Simon task as the inhibition task for about 15 minutes. During the Simon task, participants responded to either the left or right direction according to the color of the stimulus, yielding three types of trials: correctly responded trials with stimulus-response directional matching (Congruent), correctly responded trials with stimulus-response directional mismatching (Congruent), and incorrectly responded trials (Error) (Singh et al., 2023b).

To fit the fixed duration *T*_*h*_ of internal history in HSM, the dataset was downsampled to 10 Hz just before exposure to HSM. The dataset was exposed to HSM 15 times.

#### 2.4.4. Human EEG dataset recorded during resting state from patients with dementia

To study representations of brain diseases, we utilized a publicly available dataset (Miltiadous et al., 2024). This dataset was available at: https://openneuro.org/datasets/ds004504/versions/1.0.8. This EEG dataset was recorded using a 19-channel EEG system (EEG 2100, Nihon Kohden, Japan) at a sampling rate of 500 Hz during resting state. The EEG data were recorded from 29 healthy controls (HC; 11 females), 36 patients with Alzheimer’s disease (AD; 24 females), and 23 patients with frontotemporal dementia (FTD; 9 females), and the mean ages of HC, AD, and FTD were 67.90, 66.39, and 63.65 years, respectively (standard deviation = 5.40, 7.89, and 8.22). AD and FTD were determined by DSM-IIIR, DSM IV, and ICD-10. Participants remained at rest for 5.1-21.3 minutes (Miltiadous et al., 2023).

To fit the fixed duration *T*_*h*_ of internal history in HSM, the dataset was downsampled to 10 Hz just before exposure to HSM. The dataset was exposed to HSM 20 times.

#### 2.4.5. Human fMRI dataset recorded during the working memory task from patients with schizophrenia

To study representations of brain diseases, we utilized a publicly available dataset (Repovš and Barch, 2018). This dataset was available at: https://openneuro.org/datasets/ds000115/versions/00001. This fMRI dataset was recorded using a 3 T MRI system (Tim TRIO, Siemens, Germany) with a TR of 2.5 seconds during a working memory task. The fMRI data were recorded from 41 healthy controls (HC; 17 females), 23 patients with schizophrenia (SCZ; 6 females), and 35 relatives of patients with schizophrenia (rSCZ; 17 females), and the mean ages of HC, AD, and FTD were 21.03, 24.26, and 24.22 years, respectively (standard deviation = 4.76, 3.74, and 3.63). SCZ was determined by DSM-IV Axis I Disorders. Participants performed the n-back task as the working memory task for about 16 minutes. During the n-back task, participants temporarily remembered the letter presented n trials earlier and then responded with the remembered letter, yielding three types of trials: responding to the letter from 0 previous trials (0-back), responding to the letter from 1 previous trial (1-back), and responding to the letter from 2 previous trials (2-back) (Repovš and Barch, 2012).

The dataset was exposed to HSM 240 times.

#### 2.4.6. Human fMRI dataset recorded during the listening task from patients with major depressive disorder

To study representations of brain diseases, we utilized a publicly available dataset (Lepping et al., 2018). This dataset was available at: https://openneuro.org/datasets/ds000171/versions/00001. This fMRI dataset was recorded using a 3 T MRI system (Skyra, Siemens, Germany) with a TR of 3.0 seconds during a listening task. The fMRI data were recorded from 20 healthy controls (HC; 11 females) and 19 patients with major depressive disorder (MDD; 11 females), and the mean ages of HC and MDD were 29.45 and 33.53 years, respectively (standard deviation = 11.26 and 13.72). MDD was determined by SCID-I/NP. Participants performed a musical-nonmusical task as the listening task for about 24 minutes. During the musical-nonmusical task, participants listened to sounds for 30 seconds, yielding four types of trials: trials listening to positive music (Pos. music), trials listening to negative music (Neg. music), trials listening to positive non-music (Pos. non-music), and trials listening to negative non-music (Neg. non-music) (Lepping et al., 2016).

The dataset was exposed to HSM 240 times.

### 2.5. EEG processing

We preprocessed EEG data using the MATLAB toolbox EEGLAB (https://sccn.ucsd.edu/eeglab/). To remove artifacts from scalp EEG, we first applied a bandpass filter at 1-50 Hz to the raw data using a finite impulse response (FIR) filter. Eye-blink artifacts were then removed using independent component analysis (ICA). The remaining artifacts were removed using the artifact subspace reconstruction (ASR) method.

To mitigate the volume-conduction effects on scalp EEG, we applied surface Laplacian filtering to the artifact-free EEG data using the CSD toolbox (Kayser and Tenke, 2006a; Kayser and Tenke, 2006b; Kayser, 2009). We used the standard CSD parameters, m = 4 and λ = 10^−5^, as in previous studies on EEG oscillations (Tenke et al., 2011; Sihn et al., 2023; Sihn and Kim, 2024).

To isolate theta, alpha, and beta oscillations, we applied FIR bandpass filters to the surface-Laplacian-filtered data. The passband ranges of the FIR filters were 4-7, 8-12, and 13-30 Hz for theta, alpha, and beta oscillations, respectively.

To obtain amplitude envelopes of oscillations, we applied the Hilbert transform to the bandpass-filtered theta and beta oscillations. The absolute values of the Hilbert-transformed data were taken as the amplitude envelopes.

To obtain traveling waves of oscillations, we first extracted the oscillatory phases of the oscillations. To this end, we applied the Hilbert transform to the bandpass-filtered theta and beta oscillations. The angles of the Hilbert-transformed data were taken as the oscillatory phases. Based on these phases, we measured traveling waves using the local phase gradient (LPG) method developed in our previous study (Sihn and Kim, 2024). These traveling waves were spatial three-dimensional vectors and thus circular rather than linear values. Because HSM requires linear values, we converted these traveling waves into linear values. As the first step, we computed two clusters of brain-wide patterns of traveling waves. This step consisted of the following sub-steps: (1) concatenation of the three-dimensional traveling-wave data, (2) k-means clustering of these concatenated data over the time domain, and (3) reconstruction of the three-dimensional data. As the second step, the three-dimensional vectors of all channels at each time were projected onto both clusters of brain-wide traveling-wave patterns, and the resulting values, equal to twice the number of channels, represented similarity to each cluster. These values are linear and therefore suitable for HSM.

To obtain global fields of oscillations, we first extracted amplitude envelopes of oscillations. To this end, we applied the Hilbert transform to the bandpass-filtered alpha oscillations. The absolute values of the Hilbert-transformed data were taken as the amplitude envelopes. The global fields represent the average of these amplitude envelopes. We extracted peaks and troughs of the global fields as the two analysis conditions for the “Human EEG dataset recorded during resting state from patients with dementia.”

### 2.6. fMRI processing

We preprocessed fMRI data using the MATLAB toolbox Statistical Parametric Mapping 12 (SPM12; www.fil.ion.ucl.ac.uk/spm). The preprocessing procedure consisted of the following steps: (1) realignment, (2) slice-timing correction, (3) segmentation, and (4) normalization. The realignment step was performed to correct head motion. If the head motion of a participant exceeded one voxel size, we excluded that participant’s data from the analysis.

To remove potential artifacts from the fMRI data, we applied global signal regression (GSR) to the preprocessed data according to a previous study (Fox et al., 2009).

In the present study, we analyzed fMRI data based on parcellation. To this end, we utilized the Schaefer atlas with 200 parcels and 7 networks (Schaefer et al., 2018; Yeo et al., 2011). This atlas is available at: https://github.com/brainspaces/schaefer200/tree/master. After parcellation, the fMRI signals in each parcel were averaged into one-dimensional time series. Consequently, we used 200 channels of fMRI data in the analyses.

### 2.7. Methods for main analysis

Throughout this study, we used two main strategies for analysis: decoding and similarity. The decoding analysis was performed using a linear support vector machine (SVM). The classes for classification were the data conditions. For the synthetic data, a class was defined as images that share the same movement trajectory of the hill peak, that is, one cycle of going and returning. For real neural data, classes were defined as two classes: whether a certain behavior occurred or not, except for the “Human EEG dataset recorded during resting state from patients with dementia.” For that dataset, classes were defined as two classes: whether a peak of the global field occurred or not, or whether a trough of the global field occurred or not.

The other strategy was similarity analysis. The similarity analysis was performed using the cosine similarity between unit activities in the hierarchy at different time steps and under different conditions. For all data, the different conditions correspond to the different classes used in the decoding analysis.

### 2.8. Code availability

The simulation and analysis codes are available at: https://github.com/DuhoSihn/HSM.

## 3. Results

### 3.1. Synthetic data analysis

To examine whether HSM can transform data based on their spatiotemporal features, we performed a synthetic data analysis. The synthetic data consisted of images of a unimodal hill. The peak of this hill moved vertically or horizontally and then returned, forming a movie that represents multidimensional time-series data. Note that throughout this study, all data were quantized from 0 to 1 by dividing samples into 20 bins based on their rank order in scale for each channel (Figure 2A).

**Figure 2.**
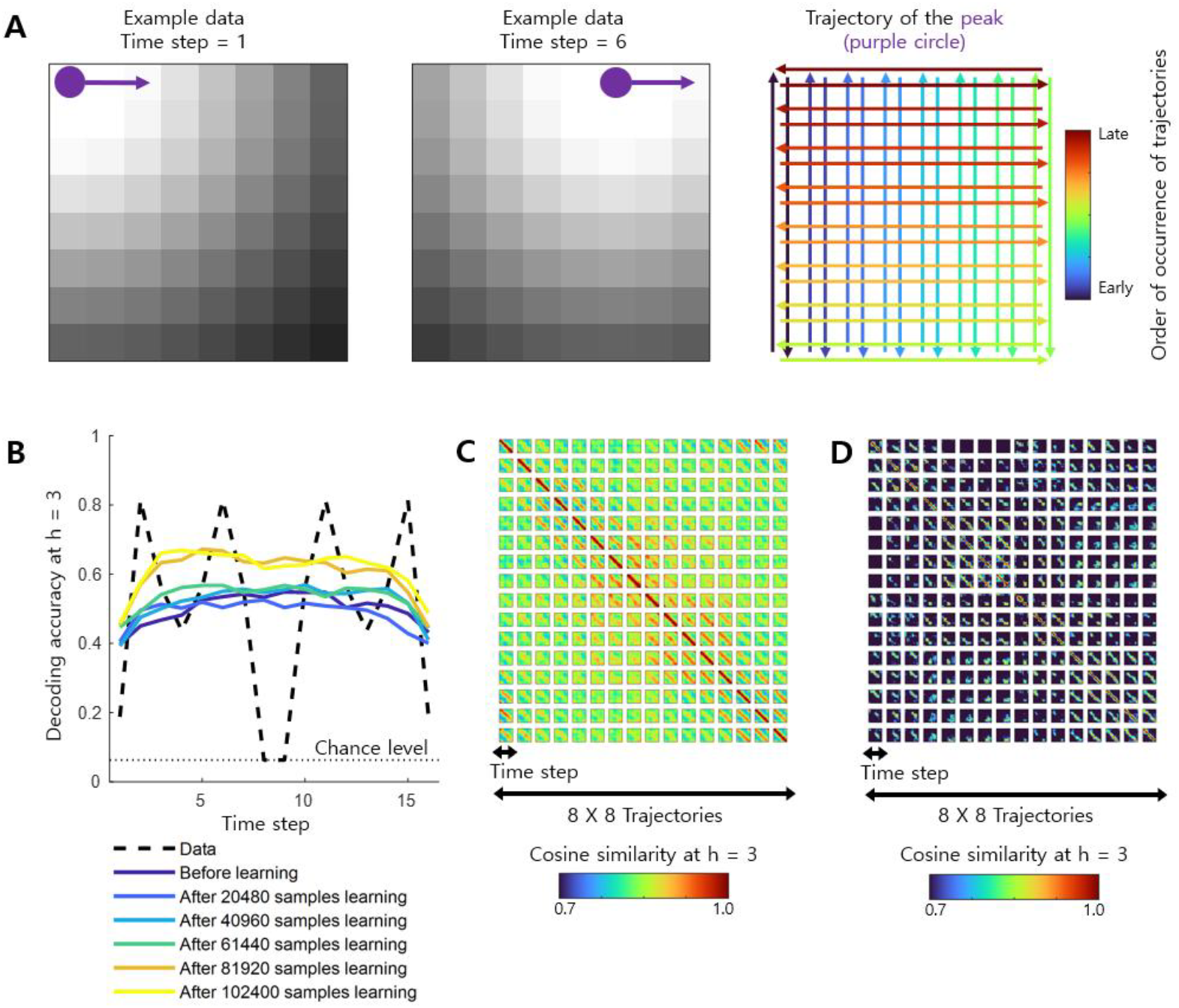
Confirmation of data transformation via HSM. (A) Description of synthetic data. The synthetic data consist of images of a unimodal hill. The peak of this hill moved vertically or horizontally and then returned, forming a movie that represents multidimensional time-series data. (B) Decoding accuracy at h = 3 for the classification of the trajectories in A. A chance level was 1/16 (16 classes). (C) Similarity result at h = 3 obtained using the cosine similarity between unit activities in the hierarchy at different time steps and for different trajectories in A. (D) Similar to C, but using the control model that has same hierarchy as HSM but is not supported history, instead of HSM.

As a result, the decoding accuracy for the synthetic data was temporally unstable, whereas the decoding accuracy became temporally stable when we used HSM. Within a class, the synthetic data consisted of images having a peak that continuously moved between very different locations, yielding temporally unstable decoding accuracies because different locations represent different time steps. In contrast, when we used HSM, it could impose the same property on continuously moving images, namely movement along a direction, through activity support from the history of past activities. The stable decoding accuracy across time steps confirmed this hypothesis, showing similar amounts of decodable information for a class despite different image locations. In other words, HSM may divide data based on trajectory rather than location, thereby exhibiting a novel spatiotemporal transformation of data. Decoding accuracy increased after learning, indicating improved HSM understanding of the synthetic data. However, an interesting point was that this stability existed even before learning, suggesting that spatiotemporal transformation could appear even before learning (Figure 2B).

The same points were observed in the similarity analysis. The beginning and end points within a class were exactly the same locations, meaning that they were the same synthetic data. However, when we used HSM, the unit activities between these two points were not similar, that is, the cosine similarity was low, indicating a spatiotemporal transformation of the data. Rather, cosine similarities between similar time steps of adjacent classes were high. This result could be interpreted to mean that information about movements at similar time steps on adjacent trajectories is mutually similar. This also verifies that HSM performs a spatiotemporal transformation of data (Figure 2C), and this was less clearly observed in the model that has same hierarchy with HSM but is not supported history (Figure 2D). These results from the similarity analysis were more evident in the upper hierarchy after learning; however, this tendency could also be observed in the lower hierarchy before learning, again suggesting that spatiotemporal transformation could appear even before learning (Figure S1).

### 3.2. Spatiotemporal transformation via HSM reveals representations of erroneous behaviors

As the next stage, we investigated the effects of HSM on the spatiotemporal transformation of real neural data. In analyses of real neural data, the decoding analysis was performed as a two-class classification of whether a certain condition (behavior) occurred or not. The similarity analysis was performed by calculating cosine similarity between unit activities in a given hierarchy at different time steps and under different conditions (behaviors).

In this context, we first analyzed various datasets in terms of erroneous behaviors. To this end, we utilized three types of publicly available datasets: human EEG data recorded during a gameplay task (naturalistic behavior), human fMRI data recorded during a working memory task, and rodent calcium-imaging data recorded during a working memory task.

We isolated theta oscillations (4-7 Hz) from the human EEG data and then extracted amplitude envelopes or measured their traveling waves. For amplitude envelopes, decoding accuracy for erroneous behavior tended to be high in the upper hierarchy, and cosine similarity between erroneous behavior and other behaviors tended to be relatively low. These results indicate that unit activities in HSM during erroneous behavior are distinct from those during other behaviors, showing the representation of erroneous behavior via spatiotemporal transformation by HSM (Figure 3A). For traveling waves, a tendency similar to that for amplitude envelopes was observed, again exhibiting the representation of erroneous behavior via spatiotemporal transformation by HSM (Figure S2A).

**Figure 3.**
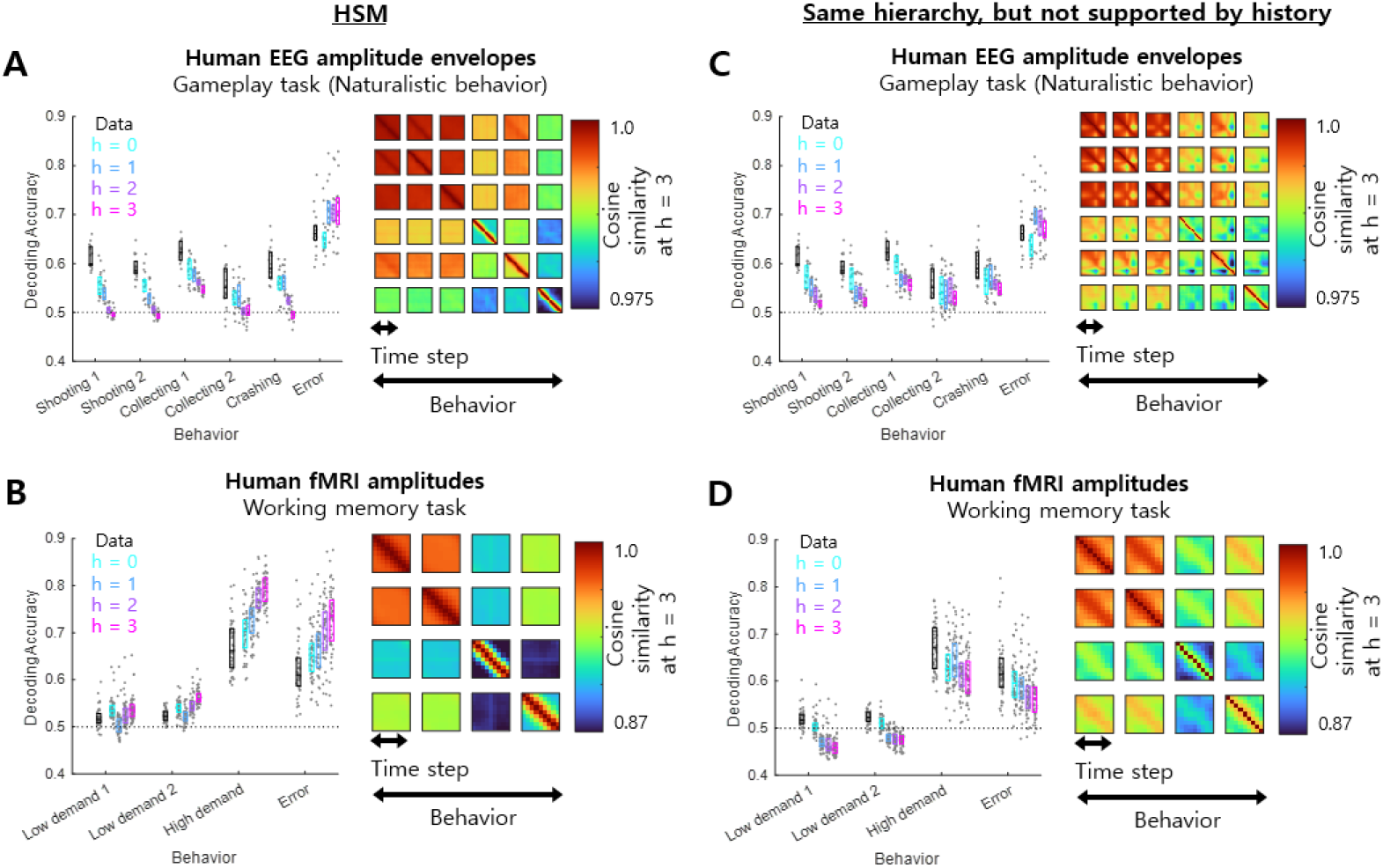
Representations of erroneous behavior via spatiotemporal transformation of HSM. **(A)** Decoding accuracies at all hierarchies and cosine similarities at h = 3 for human EEG amplitude envelopes measured during the gameplay task. (B) Decoding accuracies at all hierarchies and cosine similarities at h = 3 for human fMRI amplitudes measured during the working memory task. (C and D) Similar to A and B, respectively, but using the control model that has same hierarchy as HSM but is not supported history, instead of HSM. In all decoding-accuracy panels, each dot indicates one participant. The three horizontal lines in each box indicate the 25th, 50th, and 75th percentiles of the data.

For human fMRI data, we used amplitudes. Decoding accuracy for high demand/erroneous behaviors tended to be high in the upper hierarchy, and cosine similarity between high demand/erroneous behavior and other behaviors tended to be low. These results indicate that unit activities in HSM during high demand/erroneous behavior are distinct from those during other behaviors, showing the effect of spatiotemporal transformation via HSM. In this case, HSM did not solely isolate erroneous behavior. However, the cosine similarity between high-demand and erroneous behaviors was the lowest, indicating a strong distinction between these two behaviors. Therefore, this case could also be regarded as a representation of erroneous behavior via spatiotemporal transformation by HSM (Figure 3B).

In the control model, however, the patterns of decoding accuracy and cosine similarity shown in both EEG and fMRI datasets when HSM is used were less evident (Figure 3C and 3D), suggesting that the supporting (i.e., encoding) by history of activities is crucial for representations of erroneous behaviors.

### 3.3. Spatiotemporal transformation via HSM reveals representations of erroneous behaviors and characteristics of brain diseases

We also investigated whether these representations of erroneous behavior are present in patients with brain diseases. To this end, we analyzed human EEG data recorded during an inhibition task from healthy controls (HC) and patients with Parkinson’s disease (PD).

We isolated beta oscillations (13-30 Hz) from the human EEG data and then extracted amplitude envelopes or measured their traveling waves. For amplitude envelopes, decoding accuracy for erroneous behavior tended to be high in the upper hierarchy, and cosine similarity between erroneous behavior and other behaviors tended to be low, in both HC and PD. These results indicate that unit activities in HSM during erroneous behavior are distinct from those during other behaviors, showing the representation of erroneous behavior via spatiotemporal transformation by HSM in both HC and PD. However, decoding accuracy for erroneous behavior tended to be higher in HC than in PD, indicating a more evident representation in HC (Figure 4A). This point could also be observed in the similarity analysis. In both HC and PD, cosine similarities between erroneous behavior and other behaviors tended to be relatively low, showing the representation of erroneous behavior via spatiotemporal transformation by HSM. However, these cosine similarities were lower in HC than in PD, again indicating a more evident representation in HC. In other words, the contrast of cosine similarities between different time steps tended to be higher in HC than in PD, showing characteristics of Parkinson’s disease extracted from spatiotemporal transformation via HSM (Figure 4B). For traveling waves, a tendency similar to that for amplitude envelopes was observed, again exhibiting the representation of erroneous behavior via spatiotemporal transformation by HSM in both HC and PD, but more clearly in HC. In other words, the contrast of cosine similarities between different time steps tended to be higher in HC than in PD, showing characteristics of Parkinson’s disease extracted from spatiotemporal transformation via HSM (Figure S2B and S2C). Therefore, these results also showed representations of Parkinson’s disease.

**Figure 4.**
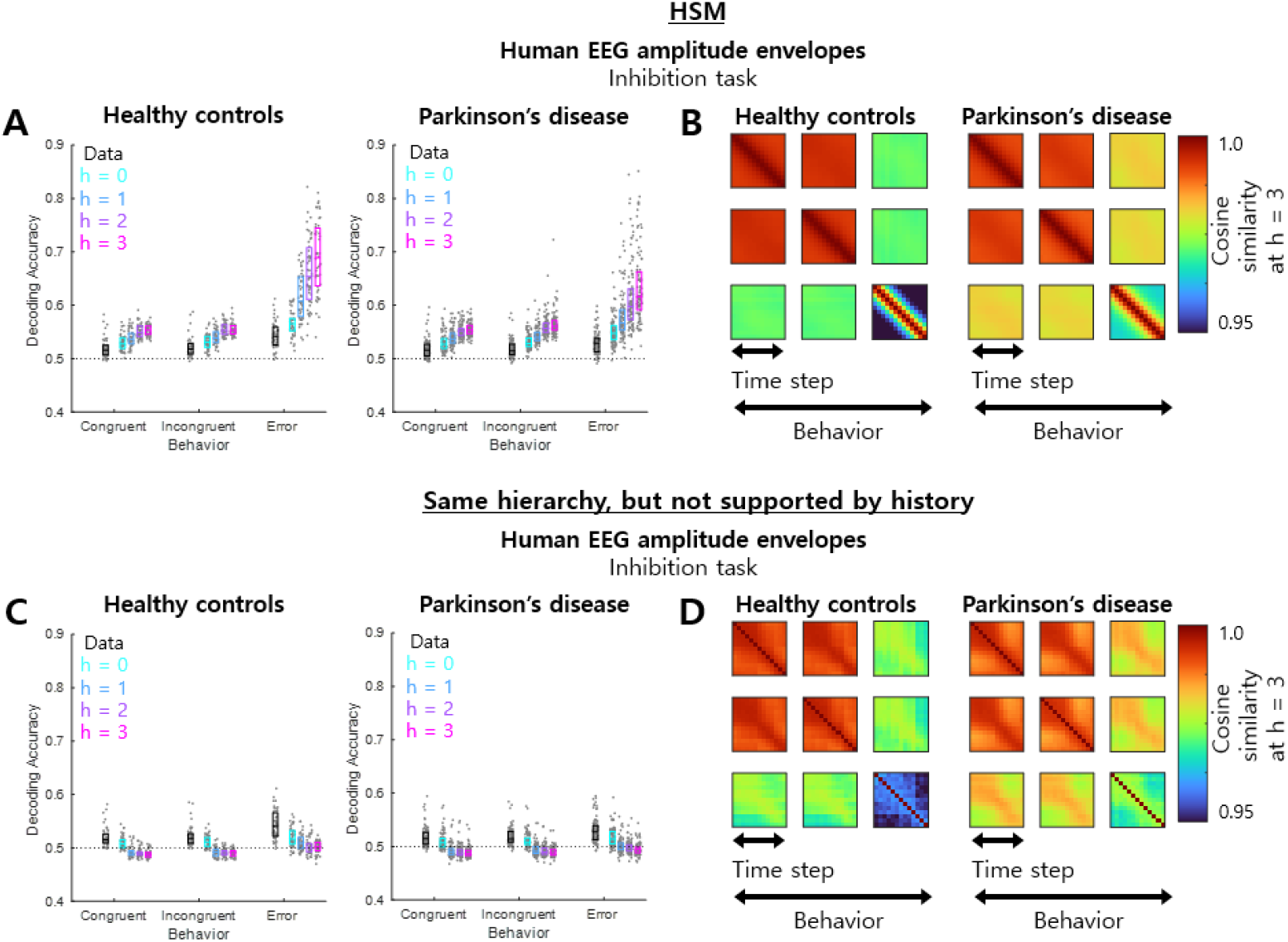
Representations of erroneous behavior and brain disease via spatiotemporal transformation of HSM. **(A)** Decoding accuracies at all hierarchies for human EEG amplitude envelopes measured during the inhibition task in healthy controls and patients with Parkinson’s disease. (B) Similar to A, but showing cosine similarities at h = 3 instead of decoding accuracies. (C and D) Similar to A and B, respectively, but using the control model that has same hierarchy as HSM but is not supported history, instead of HSM. In all decoding-accuracy panels, each dot indicates one participant. The three horizontal lines in each box indicate the 25th, 50th, and 75th percentiles of the data.

In the control model, however, the patterns of decoding accuracy in HSM were less evident for both perspectives in erroneous behavior and Parkinson’s disease (Figure 4C), suggesting that the supporting (i.e., encoding) by history of activities is crucial for representations of erroneous behavior and Parkinson’s disease. However, in cosine similarity, the HSM and the contrast model showed a similar trend (Figure 4D), which was a different result from decoding accuracy.

In addition, we explored the possibility of characteristics of other brain diseases unrelated to erroneous behavior. To this end, we utilized three types of publicly available datasets: human EEG data recorded during resting state, human fMRI data recorded during a working memory task, and human fMRI data recorded during a listening task. The human EEG data recorded during resting state were obtained from healthy controls (HC), patients with Alzheimer’s disease (AD), and patients with frontotemporal dementia (FTD). The human fMRI data recorded during the working memory task were obtained from healthy controls (HC), patients with schizophrenia (SCZ), and relatives of patients with schizophrenia (rSCZ). The human fMRI data recorded during the listening task were obtained from healthy controls (HC) and patients with major depressive disorder (MDD).

We isolated alpha oscillations (8-12 Hz) from the human EEG data, then extracted amplitude envelopes and averaged them across channels to define a global field. In this dataset, the data conditions were set to peaks or troughs of the global field. Decoding accuracies for these two conditions tended to decrease in the unit activities of HSM. The contrast of cosine similarities between different time steps tended to be higher in HC than in AD and FTD, showing characteristics of dementia extracted from spatiotemporal transformation via HSM (Figure 5A).

**Figure 5.**
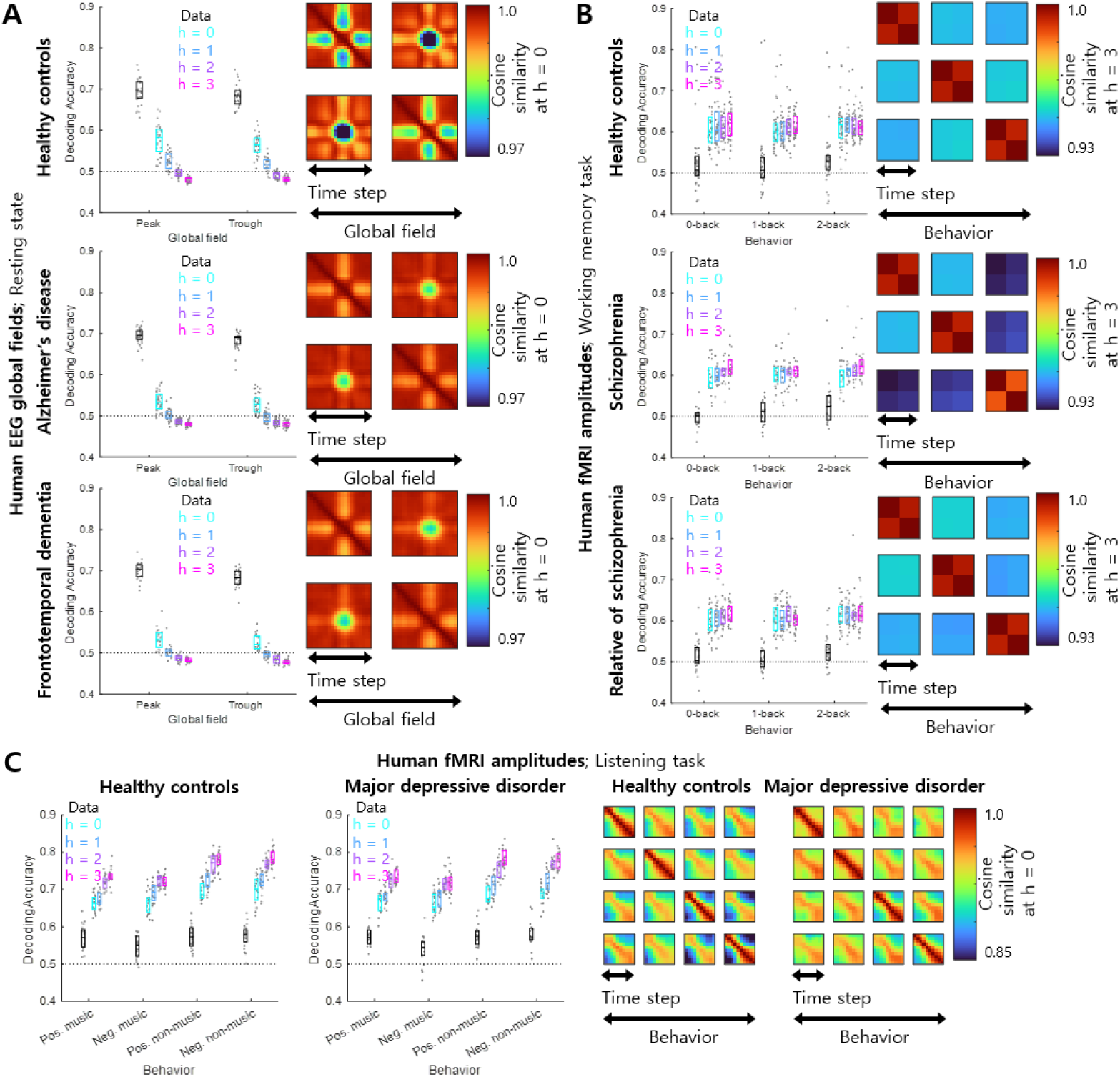
Characteristics of brain diseases via spatiotemporal transformation of HSM. (A) Decoding accuracies at all hierarchies and cosine similarities at h = 0 for human EEG global fields measured during resting state in healthy controls and patients with Alzheimer’s disease and frontotemporal dementia. (B) Decoding accuracies at all hierarchies and cosine similarities at h = 3 for human fMRI amplitudes measured during the working memory task in healthy controls, patients with schizophrenia, and relatives of patients with schizophrenia. (C) Decoding accuracies at all hierarchies and cosine similarities at h = 0 for human fMRI amplitudes measured during the listening task in healthy controls and patients with major depressive disorder. In all decoding-accuracy panels, each dot indicates one participant. The three horizontal lines in each box indicate the 25th, 50th, and 75th percentiles of the data.

For human fMRI data recorded during the working memory task, we used amplitudes. Decoding accuracies for all behaviors tended to increase in the unit activities of HSM. The contrast of cosine similarities between different behaviors tended to be lower in HC and rSCZ than in SCZ, showing characteristics of schizophrenia extracted from spatiotemporal transformation via HSM (Figure 5B).

For human fMRI data recorded during the listening task, we used amplitudes. Decoding accuracies for all behaviors tended to increase in the unit activities of HSM. The contrast of cosine similarities between different time steps tended to be higher in HC than in MDD, showing characteristics of major depressive disorder extracted from spatiotemporal transformation via HSM (Figure 5C).

### 3.4. Spatiotemporal transformation via HSM without learning

VCAF renders the variation of unit activities invariant to changes in the weight matrices at a pre-fixed level. This may lead to some degree of spatiotemporal transformation before the weight matrices are learned. It was found that spatiotemporal transformation of the synthetic data could be present even before learning (Figure 2B and S1). Therefore, we investigated the effects of learning on spatiotemporal transformation via HSM.

For representations of erroneous behavior, the trends in decoding accuracies and cosine similarities before learning were similar to those after learning (Figure S3 and S4). This consistency of trends was also observed in patients with Parkinson’s disease (Figure S4). These results indicate that representations of erroneous behavior in the present study were maintained even before learning.

For characteristics of brain disease, the trends in decoding accuracies and cosine similarities for patients with Parkinson’s disease before learning were similar to those after learning (Figure S4). However, the trends in cosine similarities for the other diseases before learning were not consistent with those after learning (Figure S5). These results indicate that characteristics of brain disease before learning in the present study were data dependent.

## 4. Discussion

In the present study, the HSM method was devised to transform multidimensional neural time-series data based on their spatiotemporal context so as to represent abnormal states such as erroneous behaviors and brain diseases. HSM spatiotemporally transforms multidimensional neural time-series data based on their spatiotemporal context, and this was confirmed through examinations of synthetic data using decoding and similarity analyses. In the HSM results, decoding accuracies were higher for erroneous behaviors than for other behaviors, and similarities were lower between erroneous behaviors and other behaviors than between other behaviors, highlighting the spatiotemporal features of erroneous behaviors. Moreover, in the HSM results, the contrast of similarities between behaviors differed between healthy controls and patients with various brain diseases, highlighting the spatiotemporal features of brain diseases. Surprisingly, many parts of these results were also present even before HSM learning, showing the virtue of HSM as a simple-to-use method.

Brain-inspired computing is expected to offer excellent performance (for reviews, see Li et al., 2024; Zhang et al., 2024; Jiao et al., 2025). It has been developed to perform better than existing systems in both software (Mou et al., 2021) and hardware (Zhang et al., 2020). Therefore, it is natural to use brain-inspired computing in the structure of HSM. In the structure of HSM, at the lower hierarchy, many modules participate simultaneously in computation. Meanwhile, at the higher hierarchy, few modules participate simultaneously in computation. This tendency can also be observed in the brain; neural activity is sparser at higher hierarchies (Vonderschen and Chacron, 2011). Furthermore. modules in higher hierarchies supported by longer past patterns of activity. This is also observed in the brain, where activity in the higher hierarchy has a longer time scale (Kiebel et al., 2008).

It has long been known that spatiotemporal features of neural data, such as EEG microstates (for a review, see Michel and Koenig, 2018) and traveling waves (for a review, see Muller et al., 2018), can represent behavioral performance. EEG microstates are related to various behavioral performances, such as mental arithmetic tasks (Kim et al., 2021), Stroop tasks (Barzon et al., 2024), audiovisual information-integration tasks (Wang et al., 2025), and naturalistic behaviors (Sihn and Kim, 2025a). Traveling waves are also related to various behavioral performances, such as visual naming tasks (Kleen et al., 2021), visual attention tasks (Alamia et al., 2023), episodic memory tasks (Mohan et al., 2024), and naturalistic behaviors (Sihn and Kim, 2025a; Sihn et al., 2026). In particular, erroneous behaviors have distinct spatiotemporal features in EEG microstates (Sihn and Kim, 2025a) and traveling waves (Alamia et al., 2023; Sihn and Kim, 2025a; Sihn et al., 2026). In the present study, decoding accuracy for erroneous behaviors in neural data was higher than that for other behaviors, although the difference was small. However, in HSM, such differences were effectively amplified, highlighting the spatiotemporal features of erroneous behaviors. This could also help reveal the mechanisms underlying erroneous behaviors.

It has long been known that spatiotemporal features of neural data, such as EEG microstates (for a review, see Michel and Koenig, 2018) and traveling waves (for a review, see Muller et al., 2018), can function as biomarkers for various brain diseases. EEG microstates are altered in patients with Parkinson’s disease (Chu et al., 2023; Meng et al., 2025; Cai et al., 2026) and dementia (Nishida et al., 2013; Teipel et al., 2021; Lian et al., 2021). Traveling waves are altered in patients with schizophrenia (Alexander et al., 2009; Castelnovo et al., 2020; Sihn and Kim, 2024) and major depressive disorder (Sihn and Kim, 2025b). For such diseases, the present study showed through similarity analysis that spatiotemporal characteristics of these diseases could be captured as altered contrasts of cosine similarities. This suggests that what previously required complex biomarkers can now be revealed through simple procedures. Moreover, decoding accuracies for erroneous behaviors in HSM subsequently differed between healthy controls and patients with Parkinson’s disease, further amplifying the difference in spatiotemporal features. These findings could help identify novel and simple biomarkers for various brain diseases.

Many aspects of the spatiotemporal transformations observed in the present study were also present even before HSM learning in analyses of both synthetic and real neural data, and this maintenance was evident in the upper hierarchy of HSM. This may be due to two reasons. First, randomized connections in HSM may produce such effects. It has been shown that randomly connected, untrained neural networks can realize many brain-like functions such as face detection (Baek et al., 2021), object-detection invariance (Cheon et al., 2022), and visual quantity comparison (Lee et al., 2023). Second, such effects of randomized connectivity would be more evident in the upper hierarchy of HSM, because lower hierarchies are directly affected by the data, thereby minimizing the effects of randomization in those hierarchies. Consequently, these results in the upper hierarchy illustrate the virtue of HSM as a simple-to-use method.

### 4.1. Limitations

In this study, HSM was learned by the Hebbian rule. Unfortunately, this learning tended to lead ultimately to trivial solutions, such as temporally unchanged unit activities. In this study, we set parameters to minimize these failures, but future research should devise methods to prevent such failures from occurring.

### 4.2. Conclusions

HSM spatiotemporally transforms multidimensional neural time-series data based on their spatiotemporal context, and this was confirmed through examinations of synthetic data using decoding and similarity analyses. In the HSM results, decoding accuracies were higher for erroneous behaviors than for other behaviors, and similarities were lower between erroneous behaviors and other behaviors than between other behaviors, highlighting the spatiotemporal features of erroneous behaviors. Surprisingly, many parts of these results were also present even before HSM learning, showing the virtue of HSM as a simple-to-use method. The HSM method can effectively highlight the spatiotemporal features of abnormal states.

## Acknowledgment

This research was supported by the National Research Foundation of Korea (NRF) grant funded by the Korea government (MSIT) (No. RS-2023-00213187) and …

## CRediT authorship contribution statement

**Duho Sihn:** Conceptualization, Methodology, Software, Validation, Formal analysis, Investigation, Resources, Data Curation, Writing - Original Draft, Writing - Review & Editing, Visualization, Supervision, Project administration, Funding acquisition. **Sung-Phil Kim:** Writing - Original Draft, Writing - Review & Editing, Visualization, Supervision, Project administration, Funding acquisition.

