## Supplementary material for "Spatiotemporal transformation of neural data reveals representations of erroneous behaviors": Figure S1; Figure S2; Figure S3; Figure S4; Figure S5

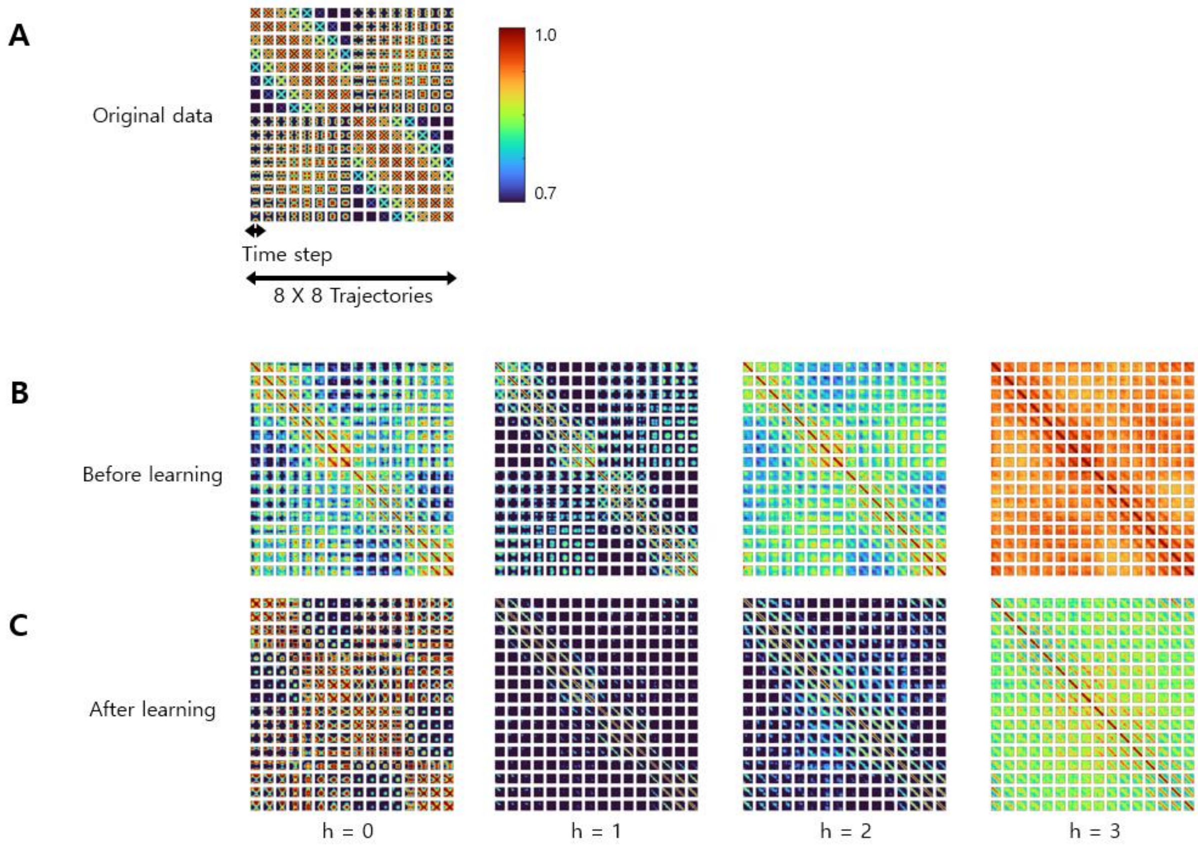


**Figure S1. Confirmation of data transformation via HSM, before and after learning.** This corresponds to Figure 2C. Synthetic data consists of images of a unimodal hill. The peak of this hill vertically or horizontally went and returned, showing the movie which represents multi-dimensional time-series data. (A) Similarity result for synthetic data obtained using the cosine similarity between unit activities of the hierarchy of different time step and different trajectories (B) Similarity result for unit activities of HSM obtained using the cosine similarity between unit activities of the hierarchy of different time step and different trajectories, before and after learning.

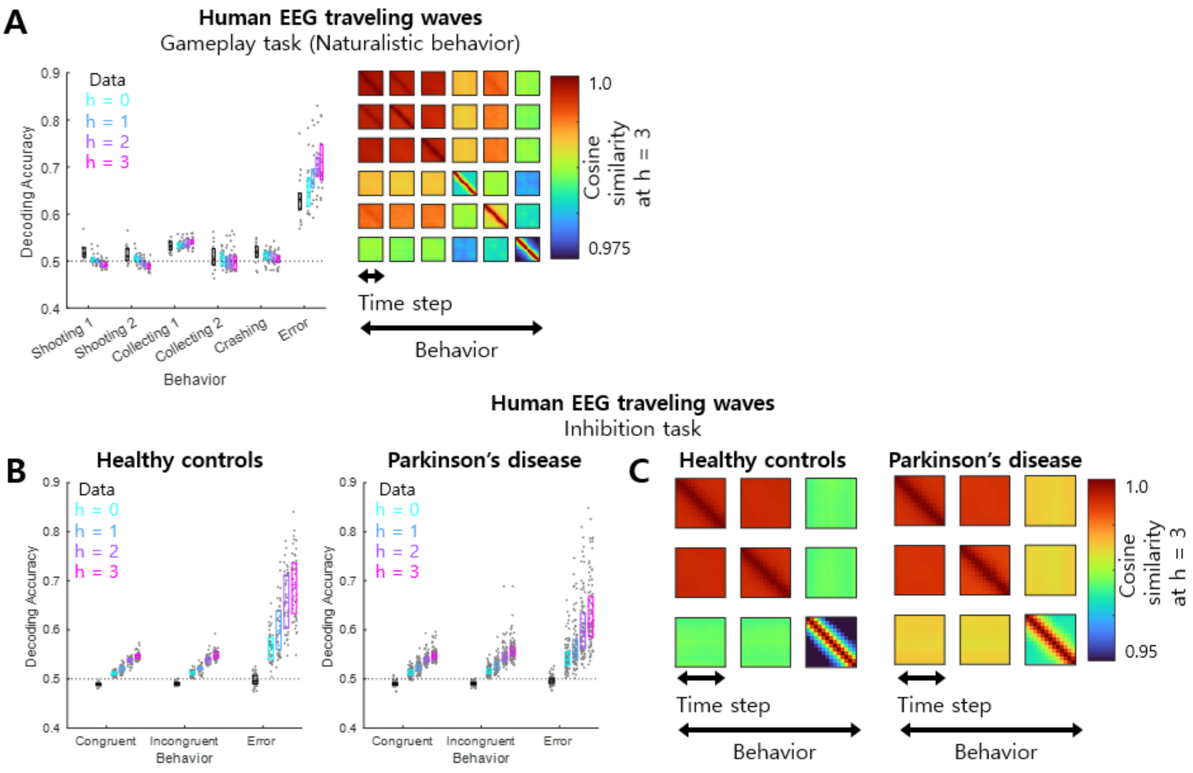


**Figure S2. Representations of erroneous behavior and brain disease via spatiotemporal transformation of HSM, in cases of traveling waves.** (A) Decoding accuracies at all hierarchies and cosine similarities at h = 3 of human EEG traveling waves measured during gameplay task. (B) Decoding accuracies at all hierarchies of human EEG traveling waves measured during inhibition task in healthy controls and patients with Parkinson's disease. (C) Similar to B, but cosine similarities at h = 3 instead of decoding accuracies.

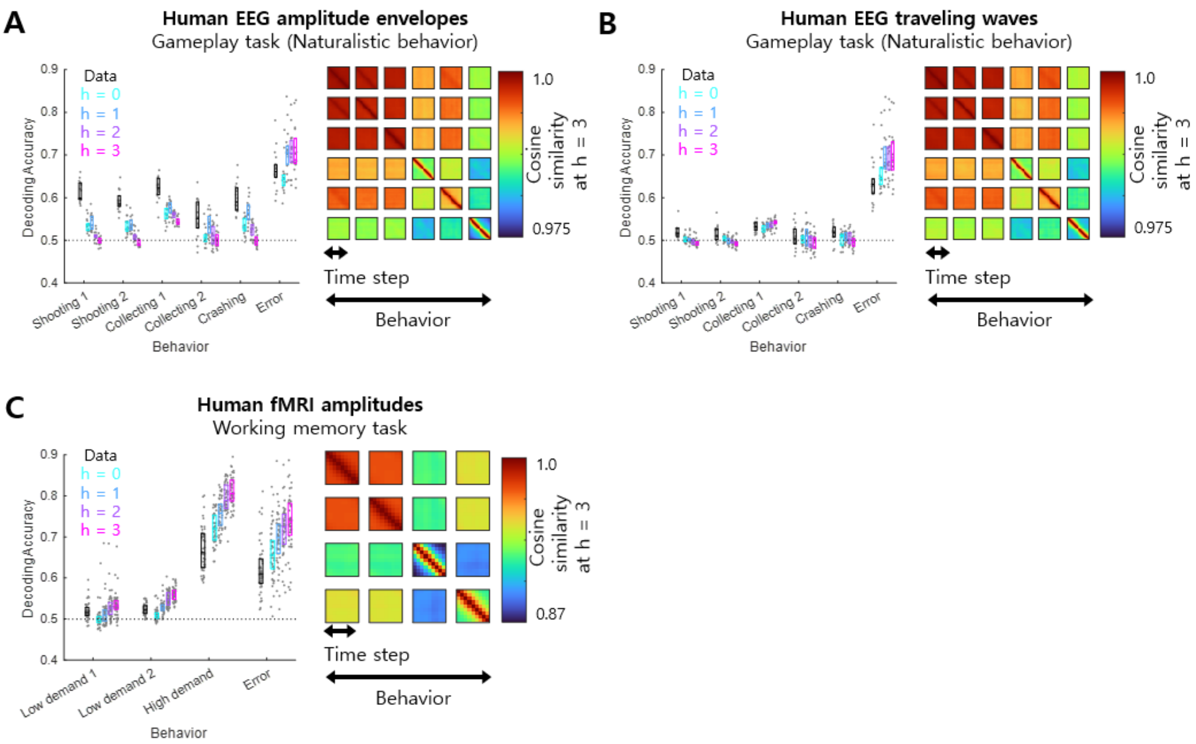


**Figure S3. Representations of erroneous behavior via spatiotemporal transformation of HSM, before learning.** This corresponds to Figure 3 and S2A. (A) Decoding accuracies at all hierarchies and cosine similarities at h = 3 of human EEG amplitudes envelopes measured during gameplay task. (B) Similar to A, but traveling waves instead of amplitude envelopes. (C) Decoding accuracies at all hierarchies and cosine similarities at h = 3 of human fMRI amplitudes measured during working memory task. In all decoding accuracy panels, each dot indicates each participant. Three horizontal lines in each box indicate 25, 50, and 75 percentiles of data.

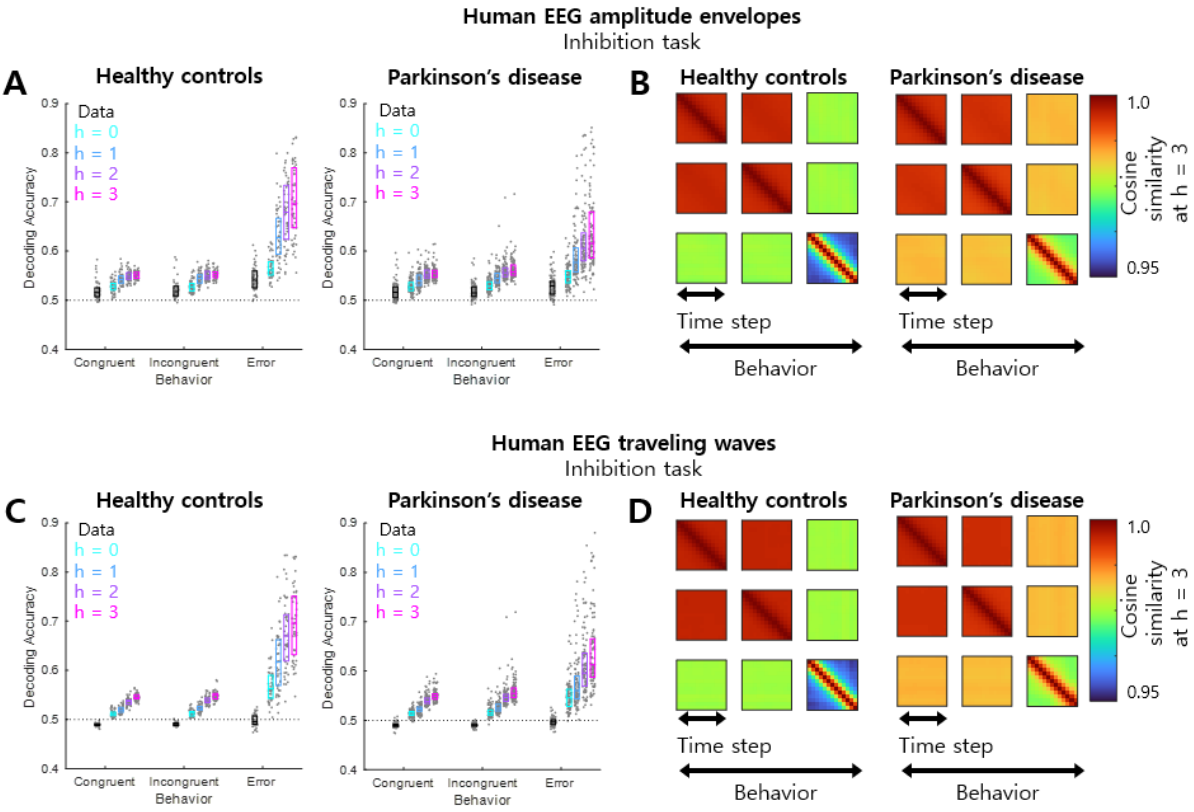


**Figure S4. Representations of erroneous behavior and brain disease via spatiotemporal transformation of HSM, before learning.** This corresponds to Figure 4, S2B, and S2C. (A) Decoding accuracies at all hierarchies of human EEG amplitudes envelopes measured during inhibition task in healthy controls and patients with Parkinson's disease. (B) Similar to A, but cosine similarities at h = 3 instead of decoding accuracies. (C) Similar to A, but traveling waves instead of amplitude envelopes. (D) Similar to C, but cosine similarities at h = 3 instead of decoding accuracies. In all decoding accuracy panels, each dot indicates each participant. Three horizontal lines in each box indicate 25, 50, and 75 percentiles of data.

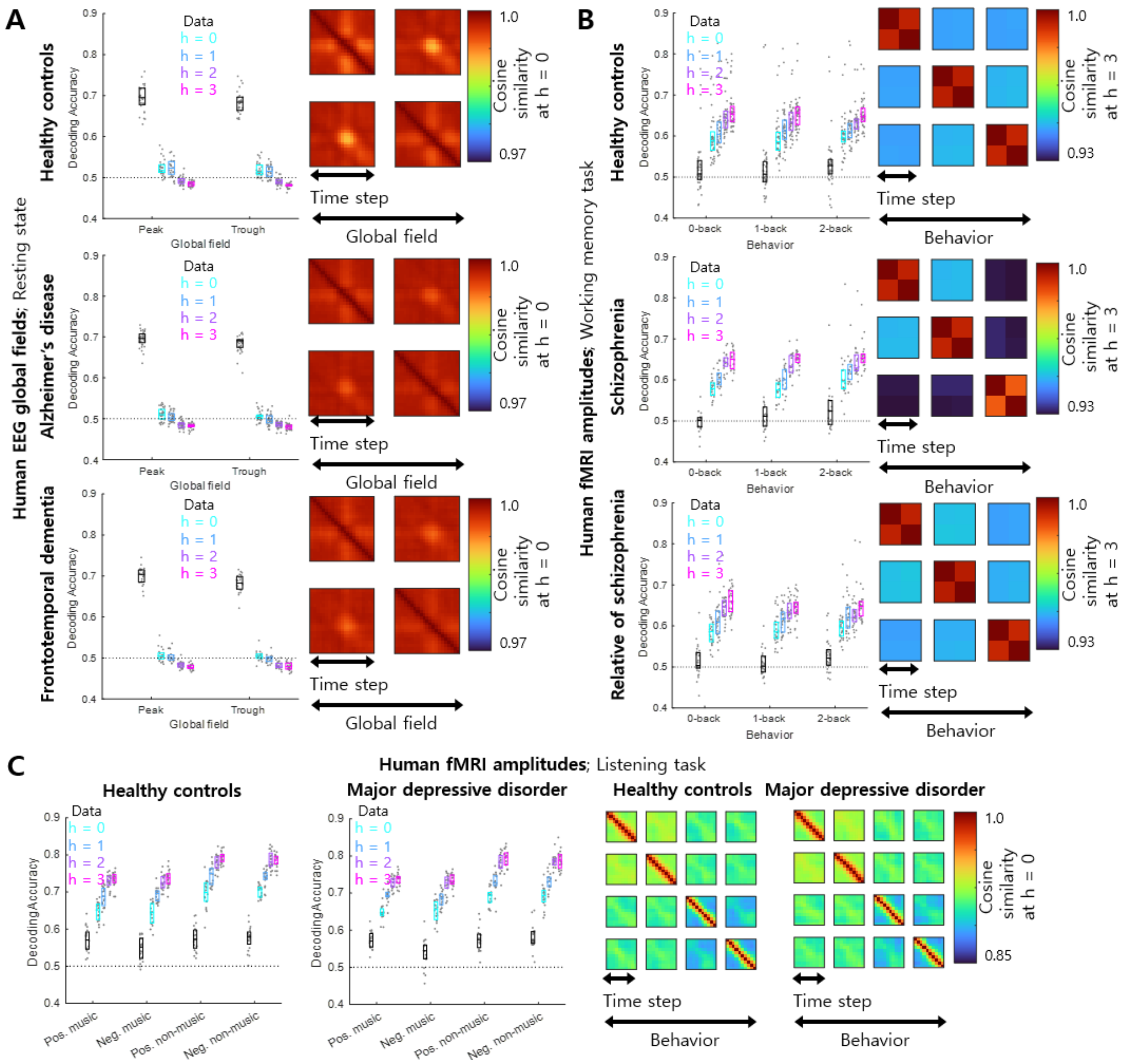


**Figure S5. Representations of brain diseases via spatiotemporal transformation of HSM, before learning.** This corresponds to Figure 5. (A) Decoding accuracies at all hierarchies and cosine similarities at h = 0 for human EEG global fields measured during resting state in healthy controls and patients with Alzheimer's disease and frontotemporal dementia. (B) Decoding accuracies at all hierarchies and cosine similarities at h = 3 for human fMRI amplitudes measured during working memory task in healthy controls, patients with schizophrenia, and relatives of patients with schizophrenia. (C) Decoding accuracies at all hierarchies and cosine similarities at h = 0 for human fMRI amplitudes measured during listening task in healthy controls and patients with major depressive disorder. In all decoding accuracy panels, each dot indicates each participant. Three horizontal lines in each box indicate 25, 50, and 75 percentiles of data.
